# Human Bocavirus Prevalence In Children With Acute Gastroenteritis From Rural Communities In The Northen Region Of South Africa

**DOI:** 10.1101/830281

**Authors:** Mpumelelo Casper Rikhotso, Ronewa Khumela, Jean Pierre Kabue, Afsatou Ndama Traoré, Natasha Potgieter

**Author notes:** Corresponding Author Mpumelelo Casper Rikhotso *, Department of Microbiology, University of Venda, Private Bag X5050, Thohoyandou, 0950, South Africa.

## Abstract

**BACKGROUND:** Acute gastroenteritis (AGE) is a leading cause of morbidity and mortality in young children worldwide. Human Bocavirus (HBoV) is an emerging virus globally associated with diarrhea. The aim of this study was to demonstrate the prevalence of HBoV genotypes in children (≤5 years) from rural communities in South Africa (SA) suffering from AGE.

**MATERIAL AND METHOD:** A total of 141 fecal samples of children ≤5 years with acute gastroenteritis (AGE) were collected from rural Primary Health Care facilities in the Vhembe district of SA between June 2017 and July 2018. Clinical symptoms and demographic data were also recorded. A total of 102 (72%) were outpatients and 39 (28%) were hospitalized patients. Human Bocavirus (HBoV) genotypes were determined using Real-Time Multiplex PCR. DNA extracts of positive samples were confirmed by conventional PCR targeting the NS1 gene. Co-infection with other enteric viruses were determined in HBoV positive samples using Real-Time PCR.

**RESULTS:** HBoV was detected in 8 (5.7%) children with AGE. Children were in the age group between 1-24 months. HBoV1 and HBoV3 genotypes were each detected in 3 (37.5%) stool samples and HBoV2 in 2 (25%) stool samples. Co-infection with other enteric viruses included Rotavirus (37.5%); Adenovirus (37.5%); Norovirus (25%) and Astrovirus (12.5%).

**CONCLUSION:** HBoV infections could be seen as a potential emerging diarrheal pathogen in South Africa. Further studies are required to understand the role of HBoV infections in children and adults with acute gastroenteritis.

**Author summary:** Acute gastroenteritis (AGE) is recognized as a major cause for mortality in children ≤5 years of age in Africa and other developing countries. Viruses known to be involved in AGE includes Rotavirus, Norovirus, Astrovirus and Adenovirus and have been reported globally. Recently the Human Bocavirus (HBoV) have been reported in numerous studies globally as a potential cause of diarrhea. In this study, the prevalence and genetic diversity of human Bocavirus in children with AGE from rural communities in Limpopo, South Africa were investigated. In total, 141 stool samples from children ≤ 5 years with AGE were assessed for the presence of HBoV using Real-Time PCR. HBoV were detected in 8 (5.7%) patients and included 3 positive samples for HBoV1 and HBoV3 respectively and 2 positive for HBoV2. No HBoV4 were detected. Among the 8 positive HBoV samples, co-infection with other enteric viruses were found in 7 (87.5%) samples, while mono infection with HBoV alone was detected in 1 (12.5%) patient. HBoV mixed infection with Rotavirus (3/8; 37.5%); Adenovirus (3/8; 37.5%); Norovirus (2/8; 25%) and Astrovirus (1/8; 12.5%) were observed in this study. This study reported for the first time on the prevalence of human Bocavirus in children with AGE from rural communities in South Africa.

## 1. Introduction

Acute gastroenteritis (AGE) is recognized as one of major causes of mortality in children ≤5 years of age in Africa and other developing countries (1, 2). AGE can be caused by several viral pathogens including Human Bocavirus (HBoV), which is an emerging viral agent reported as a potential cause of diarrhea, especially in young children (3, 4).

HBoV is a member of the *Parvoviriadae* family, *Parvovirinae* subfamily and the genus Bocavirus (4). Human Bocavirus are small, non-enveloped, icosahedral viruses’ approximately 5.3 kb single stranded DNA genome containing 3 open reading frames (ORFs): 1^st^ ORF encodes NS1, 2^nd^ ORF encodes NP1, and the 3^rd^ ORF encodes the viral proteins VP1 and VP2 (5). There are currently four Bocavirus genotypes identified globally, namely HBoV genotypes 1 to 4 (4, 6).

HBoV investigations in South Africa (SA) have only reported on the detection of HBoV genotypes in children with respiratory tract infections (7–9). There is no published work on the detection of this virus from stools in children with AGE or from children living in rural communities with poor water and sanitation infrastructure in South Africa. This study aimed to determine the prevalence and genetic diversity of HBoV in children with AGE from rural communities in Limpopo, South Africa.

## 2. Results

### 2.1 Study population characteristics

A total of 141 children with AGE were recruited in this study of which 66 (47%) were males and 75 (53%) were females. A total of 83 (59%) of the children were aged between 1-12 months, 36 (25%) of the children were aged between 13-24 months, 10 (7%) of the children were aged between 25-36 months, 9 (6%) of the children were aged between 37-48 months, and 3 (2%) of the children were aged between 49-60 months. All the children presented with symptoms of diarrhea 141 (100%), fever 48 (27%), vomiting 47 (27%), dehydration 28 (16%), respiratory tract infection 26 (15%), and abdominal pain were seen in 27 (15%) of the children.

### 2.2 Detection and genotyping of HBoV Isolates

General characteristics of HBoV positive patients is summarized in Table 2. The prevalence of HBoV genotypes 1 to 4 in stool samples of children with AGE was determined using Real-Time PCR. In total, 8 (5.7%) samples were found positive for Human Bocavirus (Table 2). HBoV1 was detected in 3 female patients with a median age of ± 9 months, HBoV2 was detected in 2 patients of which 1 was male and 1 female with a median age of ± 9.5 months, HBoV3 was detected in 3 patients (37%) of which 2 were female and 1 was male with a median age of ± 19 months (Table 2). HBoV4 was not detected in any of the 141 patients. Five (62%) of the positive cases were from clinics and 3 (37%) of the positive cases were from hospitals (Table 2).

Human Bocavirus genotypes 1 and 3 were each detected in 3 (37%) stool samples respectively, and HBoV2 was detected in 2 (25%) stool samples (Table 2) of patients. In Real-Time PCR C_T_ levels are used as a surrogate measurement of viral load in combination with standards of known quantities (10). In this study, HBoV1 was detected in 3 patients with a median C_T_ value of ± 23.8, and HBoV2 in 2 patients with a median C_T_ value of ± 36, whereas HBoV3 was detected in 3 patients with a median C_T_ value of ± 27.7 (Table 2). Genotypes were determined through Sanger DNA sequencing and confirmed by comparison with reference genotypes available in the NCBI GenBank using BLAST tool. HBoV Sequences from the current study were deposited into GenBank under accession numbers MN072357-MN072360, MN082386, MN082387.

Among the eight positive HBoV samples, single co-infection together with other enteric viruses were found in 7/8 (87.5%) patients, while mono infection with HBoV alone was detected in 1/8 (12.5%) of the HBoV positive patients. Mixed infections with Rotavirus (3/8; 37.5%); Norovirus (2/8; 25%); Adenovirus (3/8; 37.5%) and Astrovirus (1/8; 12.5%) were observed in this study population (Table 2).

General characteristics of the eight HBoV positive patient’s symptoms observed in this study. Among the 3 HBoV1 patients, one 1/3 (33.3%) had diarrhea only while two 2/3 (66.6%) had diarrhea, fever, abdominal pain, vomiting, dehydration while only 1 patient had respiratory symptoms (Table 2). In HBoV2 patients, both 2/2(100%) had diarrhea while only one 1/2 (50%) had diarrhea and fever (Table 2). All HBoV3 patients had diarrhea 3/3 (100%), from which only one 1/3 (33.3%) had diarrhea, fever, vomiting, dehydration and abdominal pain (Table 2).

HBoV 1 was observed in February (1/8; 12.5%), June (1/8; 12.5%) and July 1 (12.5%) (Figure 1). HBoV2 was only observed in June (1/8; 12.5%) and December (1/8; 12.5%), HBoV 3 was observed in March (1/8; 12.5%), July (1/8; 12.5%) and December (1/8; 12.5%) (Figure 1).

## 3. Discussion

Several studies have reported Human Bocavirus in children with respiratory tract infections in South Africa (7–9). However, no data is available on the prevalence of HBoV in children with AGE from rural communities suggesting that cases from these rural areas are most likely to be under investigated and underreported (11). This study, for the first time investigated the prevalence of HBoV in children with AGE from rural communities in South Africa.

Altogether, a total of 141 stool samples from children with AGE were assessed for the presence of HBoV. The prevalence of 5.7% for HBoV in this study was comparable to other studies which have investigated HBoV in children with diarrhea worldwide (12–14). Several of these studies have indicated that HBoV prevalence in patients with AGE ranges between 0.8% to 42% (12, 13). The high detection of HBoV in children from the clinics (outpatient) in the current study suggested that HBoV could be associated with both severe and asymptomatic diarrheal cases (15). Results from this study further suggests that HBoV could be seen as an emerging viral pathogen in the rural communities of South Africa. Although HBoV is considered a potential cause of diarrhea, available evidence supporting the causative role of the virus in acute gastroenteritis is inconclusive (16). The presence of HBoV in asymptomatic individuals raises questions regarding the role of the virus in gastrointestinal infections as a pathogen or just a bystander (16). This is due to limited available studies investigating the pathogenesis of HBoV due to the lack of cell culture system or animal model (17–19).

Reports worldwide have indicated that young children are prone to HBoV infections (20). In this study HBoV was detected in children between 1-24 months of age (≤24 months) (Table 2). No study to date have confirmed the association of HBoV infection with a specific age group of children affected. The virus could be infecting young children via fecal-oral-route transmission. A study in China (15) have reported that the transmission of HBoV was through ingestion of contaminated food/water (e.g. via flies, inadequate sanitation facilities, inadequate sewage and water treatment systems, and the cleaning food with contaminated water), direct contact with infected faces (faecal-oral-route), person-to-person contact and poor personal hygiene. In this study, the patients came from rural communities with no or inadequate water and sanitation infrastructure and poor hygiene practices (21). From the 8 positive cases of HBoV in this study, 7 (87%) used tap water as a source of water and only 1 (12%) used spring water as a source of water at home. Rural communities in South Africa usually collect water in storage tanks for both domestic and sanitation use due to scarcity of water in rural areas, this results in the contamination of water through faecal-oral-route transmission (21). Waterborne viruses represent a major health risk to population worldwide (22). There is currently limited data worldwide exploring the circulation of HBoV from environmental samples. Some studies have shown the prevalence of Human Bocavirus in river water (22, 23) and wastewater samples (24–26). Even though the role of HBoV in gastrointestinal infections still remains to be not fully understood, the risk of infection via contaminated water should be taken into consideration since many rural communities still face challenges of poor sanitation and hygiene practices (11). All 8 HBoV positive cases used a pit latrine at home. The use of a pit latrine plays a role in the transmission of pathogens since the facility lack hand washing facility and usually has flies moving in and out between the pit latrine facility and house (21).

Only, HBoV1, HBoV2 and HBoV3 were detected in this study. HBoV1 and HBoV3 were detected in 3 (37%) stool samples each, while HBoV2 was detected in 2 (25%) stool samples. Some studies have indicated that HBoV2, HBoV3 and HBoV4 are highly associated with gastroenteritis (27, 28). The widespread distribution of HBoV1 and HBoV3 in comparison to HBoV2 could presumably be due to differences in pathogenesis that may influence their transmission route and ability to establish persistence (29). A study in Thailand (30) also only detected HBoV1, HBoV2 and HBoV3. Likewise, studies from Finland and Pakistan also detected genotypes HBoV1 to 3 (31, 32). In this study we were unable to detect HBoV4. This was also the case in several other recent studies (6, 31, 33, 34), which makes the role of HBoV4 genotype unclear.

HBoV co-infection with other enteric viruses have been reported worldwide. In this study, HBoV was co-detected with other enteric viruses that are involved in acute gastroenteritis including Adenovirus F which was detected in 3/8 (37.5%) samples, while Rotavirus was detected in 3/8 (37.5%), Norovirus in 2/8 (25%) and Astrovirus in 1/8 (12.5%) samples. In only 1/8 (12.5%) of the patients, HBoV was detected alone without co-infection with other enteric virus. A study in China (33) found HBoV co-infection with Rotavirus were the most commonly detected (45.3%), followed by Human coronavirus (10.1%), Astrovirus (4.9%), and Adenovirus (4.7%). Another study in Gabon (35) co-detected HBoV with Rotavirus (33.3%), Sapovirus (33.3), and Adenovirus/Norovirus (33.3%) in children with diarrhea. In this study, Rotavirus (37.5%) and Norovirus (37.5%) were the most commonly detected, followed by, Adenovirus (25%), and astrovirus (12.5%). Previous reports have indicated that HBoV co-infection with other enteric viruses is common. Studies from China, Thailand, Japan, Brazil and Pakistan also reported co-infection were very high, while Rotavirus and Norovirus were the most predominant co-infections (13, 30, 36, 37).

A study in China (33), suggested that differences in prevalence of certain HBoV genotypes might be due to regional differences in viral epidemiology. Some studies have suggested that HBoV have a seasonal peak during the spring months, while other studies suggested the winter months, the geographic location and seasonality (38, 39) plays an important role in HBoV prevalence. In this study, samples were collected over a period of twelve months. However, samples were not available for collection in all the months. Therefore, overall HBoV prevalence was inconclusive concerning seasonal patterns.

Limitations of the study included the absence of a control group (asymptomatic cases) and mild cases of gastroenteritis. Another limitation was the small number of stool samples collected. Human Bocavirus infections are increasingly being recognized globally as a new emerging virus associated with diarrhea. Therefore, surveillance of the virus is crucial to monitor the prevalence and to help understand the role of this virus in individuals with AGE. The involvement of HBoV in children with AGE from rural communities in South Africa are most likely to be under investigated and underreported. To the best of our knowledge, this study was the first to report on the prevalence and genetic diversity of human Bocavirus in children with AGE from rural communities in Limpopo, South Africa.

## 4. Materials and methods

### 4.1 Study design

The study was a cross-sectional study and investigations were carried out between June 2017 and July 2018. Stool samples were randomly collected from patients at different Clinics (outpatients) and Hospitals (inpatients) from rural communities in the Vhembe region, Limpopo Province of South Africa. Clinics and Hospitals were both chosen because most cases of AGE in South Africa are seen by primary health care centres (Clinics) situated in the rural communities where only severe cases of AGE (with dehydration) are referred by the Clinic to the Hospital where they take over the treatment plan. In total 14 Clinics and 3 district hospitals were visited during the study period.

### 4.2 Ethical clearance and consent

The study protocol was reviewed, approved and registered by the ethics committee at the University of Venda (Ref. SMNS/17/MBY/03). Ethical clearance was obtained from the provincial Department of Health (Limpopo), South Africa (Ref: 4/2/2). Written, informed consent was given to the parent/guardian of the child to grand permission before participation and the collection of stool sample from the child.

### 4.3 Sampling

In total, 141 stool samples were collected from children ≤5 years of age with AGE, of which 102 stool specimens were collected from clinics and 39 from hospitals. Only children who fitted the criteria for acute gastroenteritis (diarrhea/vomiting/fever/cramping/dehydration) were recruited. Samples were transported to the University of Venda after collection at 4°C and stored at −20°C until further analysis.

### 4.4 Data collection

Personal information such as age and sex was collected from patients including consultation details, parental status, family living conditions, water source, and type of latrine facility used at home. Clinical symptoms, including symptoms of fever, vomiting, cough, diarrhea and dehydration were recorded.

### 4.5 DNA extraction and amplification

The Boom method was used for DNA extraction as previously described (40). The method is based on the lysing and nuclease inactivating properties of the chaotropic agent guanidinium thiocyanate, together with the nucleic acid-binding proprieties of silica particles. The extracted DNA was stored at −20ºC until further analysis.

Human Bocavirus (HBoV) Real Time PCR commercial Kit was used to detect HBoV1-4 genotypes following the manufacturer’s instructions (Liferiver ^TM^, Shanghai, China). Amplification reactions were performed in a volume of 40µl containing 35µl reaction mix, 0.4µl Enzyme Mix, 1µl Internal Control and 4µl extracted DNA according to the manufacturer’s instructions.The Real-Time m-PCR was performed using a Corbett Research Rotor Gene 6000 with the following conditions: 2 min at 37°C, 2 min at 94°C and 40 cycles of 15 sec at 93°C and 1 min 60°C. Positive samples were confirmed by sequencing.

### 4.6 Genotype analysis

All positive samples from Real-Time PCR assay were subjected to subsequent PCR targeting the NS1-nonstructural protein gene using the primers listed in Table 1.

The TopTaq Master Mix Kit (Qiagen) was used for the amplification of all positive HBoV samples following the manufactures instructions. Amplification reactions were performed in a volume of 50µl containing 25µl master mix, 5µl each of primer F and R, 15µl Free-RNase water and 5µl DNA. Amplification condition for HBoV1 were initial denaturation for 10 min at 94°C, 35 cycles of amplification (94°C for 1 min, 54°C for 1 min, and 72°C for 2 min) and the expected product size was 354 bp. HBoV2/4 amplification was initiated with a denaturation step of 15 min at 95ºC followed by 40 cycles of 94ºC for 15 sec, 53ºC for 30 sec, and 72ºC for 1 min, the expected product size was 454 bp for HBoV2/4. Human Bocavirus 2 and 4 used the same antisense primers devised by Kapoor et al. (18), which was unable to differentiate between HBoV2 and 4 genotypes. Therefore, genotypes were further confirmed through sequencing. The amplification of HBoV3 consisted of 1 step of 95ºC for 15 min followed by 45 cycles of 94ºC for 20 sec, 52ºC for 20 sec, and 72ºC for 40 sec, followed by an extension step at 72ºC for 10 min, the expected product size for HBoV3 was 440 bp (41).

All PCR products were visualized on 2% agarose gel electrophoresis and ethidium bromide staining. All positive samples were sequenced, the Sanger DNA sequencing was performed on the ABI 3500XL Genetic Analyzer POP7TM (Thermo-Scientific). The nucleotide sequences were compared with those of the reference strains available in the NCBI GenBank using BLAST tool available at http://www.ncbi.nlm.nih.gov/blast then analyzed for their respective genotypes.

### 4.7 Co-infection viruses detected

Co-infection with other enteric viruses were determined using a CFX96 (Bio-Rad) real-time PCR. The Allplex gastrointestinal panel virus assays (Seegene Technologies Inc., California, USA) was used to determine the co-detected enteric viruses following the manufactures instructions.

## 3. Acknowledgements

This study was funded by the Director of Research and innovation, University of Venda (Project number: SMNS/17/MBY/03). The National Research Foundation (NRF) is also acknowledged for funding the PhD student. The provincial and district Limpopo Departments of Health, the district executive and the nursing staff which helped to collect the clinical samples from the Vhembe district (South Africa) are acknowledged for their assistance.

